# Ethical Shades of Gray: Questionable Research Practices in Health Professions Education

**DOI:** 10.1101/256982

**Authors:** Anthony R. Artino, Erik W. Driessen, Lauren A. Maggio

## Abstract

**Purpose:** To maintain scientific integrity and engender public confidence, research must be conducted responsibly. Whereas scientific misconduct, like data fabrication, is clearly irresponsible and unethical, other behaviors—often referred to as questionable research practices (QRPs)—exploit the ethical shades of gray that color acceptable practice. This study aimed to measure the frequency of self-reported QRPs in a diverse, international sample of health professions education (HPE) researchers.

**Method:** In 2017, the authors conducted an anonymous, cross-sectional survey study. The web-based survey contained 43 QRP items that asked respondents to rate how often they had engaged in various forms of scientific misconduct. The items were adapted from two previously published surveys.

**Results:** In total, 590 HPE researchers took the survey. The mean age was 46 years (SD=11.6), and the majority of participants were from the United States (26.4%), Europe (23.2%), and Canada (15.3%). The three most frequently reported QRPs were adding authors to a paper who did not qualify for authorship (60.6%), citing articles that were not read (49.5%), and selectively citing papers to please editors or reviewers (49.4%). Additionally, respondents reported misrepresenting a participant’s words (6.7%), plagiarizing (5.5%), inappropriately modifying results (5.3%), deleting data without disclosure (3.4%), and fabricating data (2.4%). Overall, 533 (90.3%) respondents reported at least one QRP.

**Conclusions:** Notwithstanding the methodological limitations of survey research, these findings indicate that a substantial proportion of HPE researchers report a range of QRPs. In light of these results, reforms are needed to improve the credibility and integrity of the HPE research enterprise.

“Researchers should practice research responsibly. Unfortunately, some do not.” –Nicholas H. Steneck, 2006^1^

---

The responsible conduct of research is the foundation of sound scientific practice.^1,2^ The need to conduct research in a responsible manner is self-evident—if science is to inform our understanding of how the world works, it must be done in an honest, accurate, and unbiased way.^3^

Whereas behaviors like data fabrication are clearly irresponsible and highly unethical, other forms of research misconduct exploit the ethical shades of gray that color acceptable research practice. Often referred to as questionable research practices (QRPs), these behaviors “by nature of the very fact that they are often questionable as opposed to blatantly improper, also offer considerable latitude for rationalization and self-deception.”^4^ Consequently, QRPs are more prevalent and, many have argued, more damaging to science and its public reputation than obvious fraud.^4–8^ Ultimately, QRPs can waste resources, provide an unfair advantage to some researchers over others, damage the scientific record, and provide a poor example for other researchers, especially trainees.^7^

Health professions education (HPE) is not immune to the damaging effects of irresponsible research practices. In the HPE context, we define QRPs as poor data management; inappropriate research procedures, including questionable procedures for obtaining informed consent; insufficient respect and care for study participants; improper research design; carelessness in observation and analysis; suboptimal trainee and mentor partnerships; unsuitable authorship or publishing practices; and derelictions in reviewing and editing.^7^ The need to guard against such practices is frequently described in the author instructions for most scientific journals. For example, *Academic Medicine*’s author instructions clearly describe ethical considerations related to authorship, prior and duplicate publications, conflicts of interests, and ethical treatment of human subjects (http://journals.lww.com/academicmedicine/_layouts/15/1033/oaks.journals/informationforauthors.aspx). Such journal guidelines are often patterned on the recommendations of the Committee on Publication Ethics (https://publicationethics.org/).

In the last decade, commentaries by several HPE journal editors have highlighted instances of QRPs in article submissions, including self-plagiarism, so-called “salami slicing” (i.e., inappropriately dividing a single study into multiple papers), and unethical authorship practices.^9–11^ Additionally, a recent review of four HPE journals found that 13% of original research articles published in 2013 did not address approval by an ethics review board or stated that it was unnecessary, without further discussion.^12^ Moreover, a 2017 study of senior HPE leaders highlighted multiple problematic authorship practices, including honorary authorship and the exclusion of authors who deserved authorship.^13^ Notably, only about half of the senior researchers surveyed were able to correctly identify the authorship standards used by most medical journals (i.e., the International Committee of Medical Journal Editors (ICMJE) authorship criteria^14^).

Notwithstanding these examples, QRPs have received limited attention in the HPE literature. In a recent article,^7^ we attempted to raise the community’s awareness of QRPs and highlight the need to examine their pervasiveness among HPE researchers. With this call to action in mind, we conducted the present study to measure the frequency of self-reported QRPs in a diverse, international sample of HPE researchers. In doing so, we hope to continue the conversation about QRPs in our growing HPE field, with the ultimate goal of promoting the responsible conduct of high-quality, ethical research.

## Method

To measure the frequency of serious research misconduct and other QRPs, many different approaches have been employed in the literature. These include counts of confirmed cases of researcher fraud and paper retractions, as well research audits by government funders.^8^ Such methods are limited because they are calculated based on misconduct that has been discovered, and detecting such misconduct is difficult.^15^ Moreover, distinguishing intentional misconduct from honest mistakes is challenging. Therefore, such approaches significantly underestimate the real frequency of QRPs, since only researchers know if they have willfully acted in a questionable or unethical manner.

To address these challenges, survey methods have been used to ask scientists directly about their research behaviors.^4–6,16,17^ Like the measurement of any socially undesirable behavior, assessing QRPs via self-report likely underestimates the true prevalence or frequency of the behaviors. Nonetheless, when employed appropriately, survey methods can generate reasonable estimates that provide a general sense of the problem’s scope.^18,19^

Therefore, we administered an anonymous, cross-sectional survey to determine the frequency of QRPs in a sample of HPE researchers. The Ethical Review Board Committee of the Netherlands Association for Medical Education approved this study (Dossier #937).

## Survey development

We developed our survey instrument by adapting several existing surveys. The final version of the survey featured a total of 66 items divided into three sections (see Supplemental Digital Appendix). The first section included 43 items derived from two previously published surveys assessing QRPs in biomedicine.^5,6^ The QRP items asked respondents to identify how often they had engaged in the particular research practice. The practices spanned the research continuum, from data collection and storage to study reporting, collaboration, and authorship. The QRP items employed a six-point, Likert-type, frequency-response scale: *never*, *once*, *occasionally*, *sometimes*, *frequently*, and *almost always*. Each item also included the response option *not applicable to my work*.

We slightly modified the original QRP items to improve their clarity and relevance to the HPE research context. For example, the original item “inadequately handled or stored data or (bio)materials” was revised to “inappropriately stored sensitive research data (e.g., data that contains personally identifiable information).” Following these modifications, 19 experienced HPE researchers reviewed the adapted survey items and provided detailed qualitative feedback.^20^ Ten of the expert reviewers were women, and all held doctoral degrees (13 PhDs, 4 MD/PhDs, and 2 MDs). On average, the reviewers had published 98.9 journal articles (SD=66.1) findable in PubMed. Based on the date of their first HPE publication, they had been publishing in the field for an average of 20.7 years (SD=9.3). Expert reviewers reported their work location as the United States (n=9), Canada (n=3), Europe (n=2), South America (n=1), Africa (n=1), and Australia (n=1), and all but two were identified as full professors.

The expert feedback included comments on item relevance, clarity, missing facets, and suggestions for overall survey improvement. Based on the expert feedback, we revised the survey again; revisions included wording modifications and the development of several new items specific to HPE research. For example, based on the recommendations of several experts, we created the following items related to qualitative research methods, among several others: “misrepresented a participant’s words or writings” and “claimed you used a particular qualitative research approach appropriately (e.g., grounded theory) when you knowingly did not.”

The second section of the survey included nine publication pressure items^16^ (these data are not reported here), and the final section included 13 demographic items. The survey ended with a single, open-ended question, which gave respondents the opportunity to give feedback on the survey instrument itself or provide additional thoughts on QRPs in HPE.

## Sampling and survey distribution procedures

To create our sample, we used two separate approaches. First, we created a “curated sample” by searching Web of Science, Scielo (a database focused on South America), African Journals Online, and Asia Journals Online for articles in over 20 HPE journals published in 2015 and 2016. From these articles, we extracted all available author email addresses, removing duplicate authors. This process generated a sample of 1,840 unique HPE researchers. All names and emails were entered into Qualtrics, an online survey tool (Qualtrics, Provo, Utah), and the survey was then distributed in four waves of email invitations: wave 1 (sent November 13, 2017), wave 2 (sent November 20, 2017), wave 3 (sent November 27, 2017), and wave 4 (sent December 11, 2017).

Next, we collected a “social media sample” by posting anonymous links to the survey on our Twitter and Facebook accounts (posted on December 11, 2017). All survey responses obtained from the social media links were tracked separately from those sent to the curated sample. To prevent duplicate submissions, respondents in the social media sample were given the option to select “I have already completed this survey” on the informed consent page.

## Statistical analyses

Prior to analysis, we screened the data for accuracy and missing values. Next, we calculated the response rate in the curated sample using response rate definition #6, as delineated by the American Association for Public Opinion Research.^21^ Then, to assess potential nonresponse bias in the curated sample, we used wave analysis to calculate a nonresponse bias statistic.^22^ In wave analysis, late respondents are considered proxies for non-responders, and their responses are compared to responses from the first wave. In addition, to determine whether or not it was appropriate to combine the curated and social media samples for analysis, we conducted a multivariate analysis of variance (MANOVA) to compare respondents on several demographic characteristics: age, experience doing HPE research, percentage of work time dedicated to HPE research, and number of peer-reviewed publications. Finally, we calculated descriptive statistics for the total sample, with particular emphasis on the frequency of self-reported QRPs. All data analyses and visualizations were conducted using IBM SPSS Statistics (IBM Corporation, New York, NY) and Microsoft Excel (Microsoft Corporation, Redmond, WA), respectively.

## Results

Of the 1,840 email invitations sent to HPE researchers in the curated sample, 199 were returned as undeliverable, leaving 1,641 potential respondents. Of these, 463 (28.2%) researchers completed a least a portion of the survey.^21^ Results from the wave analysis revealed a nonresponse bias statistic of 0.36. On a six-point, frequency-response scale, this represents a 6% difference, which is unlikely to have a meaningful effect on practical interpretation of the results.^22^

The social media sample yielded an additional 127 responses. Results from the MANOVA comparing respondents in the curated sample to those in the social media sample revealed statistically significant differences between the two groups F(5, 524) = 6.67, P < .001. In particular, post-hoc analyses revealed that respondents in the social media sample were slightly younger (M=40.7 years) and more inexperienced in HPE research (M=7.5 years) than those in the curated sample (M=47.4 years and M=11.0 years, respectively). That said, the two groups did not differ in terms of percentage of work time spent doing HPE research activities or mean number of peer-reviewed publications. Therefore, because our goal was to explore the frequency of QRPs among a diverse, international sample of HPE researchers, we elected to pool the two samples and analyze the data together.

Of the 590 respondents in the pooled sample, the mean age was 46 years (SD=11.6), and there were 305 (51.7%) women, 246 (41.7%) men, and 39 (6.6%) individuals who did not report their gender. As indicated in Table 1, the sample consisted of HPE researchers from across the World Health Organization’s six world regions. The majority reported their location as the United States (26.4%), Europe (23.2%), and Canada (15.3%). Respondents’ education, area of study, work context and role, academic rank, and primary research activities are also presented in Table 1. In addition, respondents reported the following: years involved in HPE in any capacity (M=14.9 years, SD=9.7), years involved in HPE research (M=11.3 years, SD=8.5), percentage of work time spent conducting HPE research (M=27.3%, SD=23.7%), and total number of peer-reviewed publications (M=40.1, SD=55.0).

**Table 1.** Demographic characteristics of an international sample of 590 health professions education researchers.

| Demographic characteristic |  | No. (%) |
| --- | --- | --- |
| Gender | Female | 305 (51.7) |
|  | Male | 246 (41.7) |
|  | Not reported | 39 (6.6) |
| Region | United States | 156 (26.4) |
|  | Europe (not including United Kingdom) | 96 (16.3) |
|  | Canada | 90 (15.3) |
|  | Australia and New Zealand | 42 (7.1) |
|  | United Kingdom | 41 (6.9) |
|  | Africa | 37 (6.3) |
|  | Asia | 37 (6.3) |
|  | South/Latin America and Caribbean | 18 (3.0) |
|  | Middle East | 16 (2.7) |
|  | Other | 12 (2.0) |
|  | Not reported | 45 (7.6) |
| Education <sup>a</sup> | Bachelor's degree | 245 (44.4) |
|  | Master's degree | 306 (55.5) |
|  | MD/DO degree | 276 (50.0) |
|  | PhD/EdD degree | 323 (58.5) |
|  | Other professional degree | 30 (5.4) |
| Area of study for highest degree selected | Social science | 293 (49.7) |
|  | Clinical science | 167 (28.3) |
|  | Other | 42 (7.1) |
|  | Basic science | 39 (6.6) |
|  | Humanities | 11 (1.9) |
|  | Not reported | 38 (6.4) |
| Work context <sup>a</sup> | Undergraduate medical education | 353 (65.7) |
|  | Graduate medical education | 308 (57.4) |
|  | Continuing medical education | 142 (26.4) |
| Work role | Researcher | 174 (29.5) |
|  | Clinician | 136 (23.1) |
|  | Administrator/program director | 89 (15.1) |
|  | Teacher | 86 (14.6) |
|  | Other | 65 (11.0) |
|  | Not reported | 40 (6.8) |
| Academic rank | Professor | 140 (23.7) |
|  | Associate professor | 109 (18.5) |
|  | Assistant professor | 99 (16.8) |
|  | Lecturer/instructor | 64 (10.9) |
|  | Graduate student | 42 (7.1) |
|  | Postdoctoral fellow | 23 (3.9) |
|  | Resident | 16 (2.7) |
|  | Other | 52 (8.8) |
|  | Not reported | 45 (7.6) |
| Primary research activity | Mixed-method | 280 (47.5) |
|  | Quantitative | 149 (25.3) |
|  | Qualitative | 119 (20.2) |
|  | Not reported | 42 (7.1) |

Abbreviations: MD, Doctor of Medicine; DO, Doctor of Osteopathic Medicine.
<sup>a</sup> Percentages total more than 100% because respondents could select more than one category.

Table 2 summarizes the frequency of self-reported QRPs, and the Figure provides a visual representation of these results. To simplify the figure, we collapsed the response options of *occasionally*, *sometimes*, *frequently*, and *almost always* into a single frequency option labeled *more than once*. Finally, the Box lists the top 10 most frequently reported QRPs among our respondents, from highest to lowest frequency. As indicated, the most frequently reported QRPs were related to authorship and study reporting practices, as well as issues around data storage, collection, and interpretation. Additionally, 39 (6.7%) respondents reported misrepresenting a participant’s words, 31 (5.5%) reported using sections of text from another author’s copyrighted material without permission or proper citation, 30 (5.3%)reported inappropriately modifying study results due to pressure from a research advisor or collaborator, 20 (3.4%) reported deleting data before performing analysis without disclosure, and 14 (2.4%) reported fabricating data. Overall, 533 (90.3%) respondents reported at least one QRP.

**Table 2.** Freqquency of self-reported questionable research practices among an international sample of 590 health fessions education researchers, reported as No. (%)^a^.

| Data Collection and Storage |  |  |  |  |  |  |  |  |
| --- | --- | --- | --- | --- | --- | --- | --- | --- |
|  | N | Never | Once | Occasionally | Sometimes | Frequently | Almost always | Not applicable |
| 1-Conducted a human-subjects research study without ethics approval (i.e., without institutional review board [IRB] approval) | 589 | 465 (78.9) | 50 (8.5) | 38 (6.5) | 13 (2.2) | 6 (1.0) | 4 (0.7) | 13 (2.2) |
| 2-Circumvented one or more aspects of human-subjects ethics rules (i.e., IRB rules) | 585 | 463 (79.1) | 47 (8.0) | 49 (8.4) | 10 (1.7) | 4 (0.7) | 0 (0.0) | 12 (2.1) |
| 3-Collected course or curriculum data under the guise of "program evaluation" without human-subjects ethics (IRB) approval with the ultimate intent of using the data for research purposes | 588 | 420 (71.4) | 45 (7.7) | 65 (11.1) | 24 (4.1) | 9 (1.5) | 2 (0.3) | 23 (3.9) |
| 4-Inappropriately stored sensitive research data (e.g., data that contains personally identifiable information) | 585 | 353 (60.3) | 56 (9.6) | 119 (20.3) | 36 (6.2) | 15 (2.6) | 2 (0.3) | 4 (0.7) |
| 5-Inappropriately emailed sensitive research data (e.g., data that contains personally identifiable information) | 585 | 436 (74.5) | 36 (6.2) | 84 (14.4) | 20 (3.4) | 3 (0.5) | 0 (0.0) | 6 (1.0) |
| 6-Stopped collecting data earlier than planned because the results already reached statistical significance, without formal stopping rules | 586 | 497 (84.8) | 22 (3.8) | 13 (2.2) | 6 (1.0) | 3 (0.5) | 1 (0.2) | 44 (7.5) |
| 7-Fabricated data | 586 | 570 (97.3) | 10 (1.7) | 2 (0.3) | 1 (0.2) | 0 (0.0) | 1 (0.2) | 2 (0.3) |
| 8-Pressured a student or other subordinate to be a study participant in your research | 583 | 536 (91.9) | 14 (2.4) | 20 (3.4) | 4 (0.7) | 1 (0.2) | 0 (0.0) | 8 (1.4) |
| 9-Used students or residents as research subjects without informing the overseeing dean, program director, or other pertinent official | 584 | 468 (80.1) | 23 (3.9) | 46 (7.9) | 17 (2.9) | 8 (1.4) | 2 (0.3) | 20 (3.4) |
| Data Analysis |  |  |  |  |  |  |  |  |
|  | N | Never | Once | Occasionally | Sometimes | Frequently | Almost always | Not applicable |
| 10-Deleted data before performing data analysis without disclosure | 582 | 553 (95.0) | 9 (1.5) | 11 (1.9) | 0 (0.0) | 0 (0.0) | 0 (0.0) | 9 (1.5) |
| 11-Ignored a colleague's use of flawed data | 581 | 447 (76.9) | 36 (6.2) | 46 (7.9) | 16 (2.8) | 5 (0.9) | 1 (0.2) | 30 (5.2) |
| 12-Ignored a colleague's questionable interpretation of data | 581 | 404 (69.5) | 44 (7.6) | 88 (15.1) | 17 (2.9) | 2 (0.3) | 2 (0.3) | 24 (4.1) |
| 13-Reported a downwardly rounded p-value (e.g., reporting that a p-value of .054 is less than .05) | 581 | 513 (88.3) | 11 (1.9) | 8 (1.4) | 0 (0.0) | 0 (0.0) | 0 (0.0) | 49 (8.4) |
| 14-Misrepresented a participant's words or writings | 580 | 526 (90.7) | 16 (2.8) | 21 (3.6) | 2 (0.3) | 0 (0.0) | 0 (0.0) | 15 (2.6) |
| 15-Decided whether to exclude non-outlier data after looking at the impact of doing so on the results | 584 | 470 (80.5) | 32 (5.5) | 32 (5.5) | 7 (1.2) | 1 (0.2) | 0 (0.0) | 42 (7.2) |
| 16-In a qualitative study, failed to report disconfirming examples or cases that weaken your conclusions | 583 | 422 (72.4) | 32 (5.5) | 32 (5.5) | 6 (1.0) | 1 (0.2) | 0 (0.0) | 90 (15.4) |
| 17-Collected more data after seeing that the results were almost statistically significant | 583 | 434 (74.4) | 33 (5.7) | 40 (6.9) | 4 (0.7) | 3 (0.5) | 1 (0.2) | 68 (11.7) |
| 18-To confirm a hypothesis, selectively deleted or changed data after performing data analysis | 583 | 523 (89.7) | 16 (2.7) | 6 (1.0) | 0 (0.0) | 0 (0.0) | 0 (0.0) | 38 (6.5) |
| 19-Reported an unexpected finding as having been hypothesized from the start | 583 | 433 (74.3) | 47 (8.1) | 55 (9.4) | 9 (1.5) | 4 (0.7) | 2 (0.3) | 33 (5.7) |
| 20-Concealed results that contradicted your previous findings or convictions | 583 | 538 (92.3) | 23 (3.9) | 11 (1.9) | 1 (0.2) | 0 (0.0) | 0 (0.0) | 10 (1.7) |
| 21-Claimed you used a particular qualitative research approach appropriately (e.g., grounded theory) when you knowingly did not | 566 | 432 (76.3) | 39 (6.9) | 22 (3.9) | 3 (0.5) | 0 (0.0) | 0 (0.0) | 70 (12.4) |
| 22-Claimed you used a particular qualitative research technique appropriately (e.g., saturation, triangulation) when you knowingly did not | 566 | 433 (76.5) | 42 (7.4) | 18 (3.2) | 3 (0.5) | 0 (0) | 0 (0) | 70 (12.4) |
| **Study Reporting** |  |  |  |  |  |  |  |  |
|  | N | Never | Once | Occasionally | Sometimes | Frequently | Almost always | Not applicable |
| 23-Spread study results over more papers than is appropriate (so-called "salami slicing") | 566 | 442 (78.1) | 59 (10.4) | 52 (9.2) | 2 (0.4) | 1 (0.2) | 0 (0.0) | 10 (1.8) |
| 24-Deliberately failed to mention important limitations of a study in the published paper | 566 | 517 (91.3) | 19 (3.4) | 26 (4.6) | 3 (0.5) | 0 (0.0) | 0 (0.0) | 1 (0.2) |
| 25-Deliberately failed to mention an organization that funded your research in the published paper | 565 | 549 (97.2) | 5 (0.9) | 1 (0.2) | 0 (0.0) | 0 (0.0) | 0 (0.0) | 10 (1.8) |
| 26-Inappropriately modified the results of a study due to pressure from a research advisor or other collaborator | 564 | 531 (94.1) | 21 (3.7) | 8 (1.4) | 1 (0.2) | 0 (0.0) | 0 (0.0) | 3 (0.5) |
| 27-Inappropriately modified the results of a study due to pressure from a funding agency | 566 | 532 (94.0) | 5 (0.9) | 0 (0.0) | 0 (0.0) | 0 (0.0) | 0 (0.0) | 29 (5.1) |
| 28-Failed to disclose relevant financial or intellectual conflicts of interest | 567 | 548 (96.6) | 6 (1.1) | 4 (0.7) | 1 (0.2) | 0 (0.0) | 0 (0.0) | 8 (1.4) |
| 29-Used someone else's ideas without their permission or proper citation | 567 | 536 (94.5) | 15 (2.6) | 14 (2.5) | 1 (0.2) | 0 (0.0) | 0 (0.0) | 1 (0.2) |
| 30-Used sections of text from another author's copyrighted material without permission or proper citation | 566 | 534 (94.3) | 13 (2.3) | 16 (2.8) | 2 (0.4) | 0 (0.0) | 0 (0.0) | 1 (0.2) |
| 31-Used sections of text from your own publications without proper citation (so-called "self-plagiarism") | 564 | 449 (79.6) | 35 (6.2) | 65 (11.5) | 12 (2.1) | 0 (0.0) | 0 (0.0) | 3 (0.5) |
| 32-Selectively cited certain papers just to please editors or reviewers | 566 | 284 (50.2) | 57 (10.1) | 170 (30.0) | 39 (6.9) | 7 (1.2) | 7 (1.2) | 2 (0.4) |
| 33-Cited articles and or materials that you have not read | 565 | 284 (50.3) | 39 (6.9) | 181 (32.0) | 48 (8.5) | 12 (2.1) | 0 (0.0) | 1 (0.2) |
| 34-Selectively cited your own work just to improve your citation metrics | 562 | 391 (69.6) | 28 (5.0) | 101 (18.0) | 24 (4.3) | 15 (2.7) | 2 (0.4) | 1 (0.2) |
| 35-Reused previously published data without disclosure (co-called "duplicate publication") | 564 | 545 (96.6) | 11 (2.0) | 4 (0.7) | 1 (0.2) | 0 (0.0) | 0 (0.0) | 3 (0.5) |
| 36-Used confidential information obtained as a reviewer or editor for your own research or publications | 564 | 543 (96.3) | 11 (2.0) | 6 (1.1) | 0 (0.0) | 0 (0.0) | 0 (0.0) | 4 (0.7) |
| **Collaboration and Authorship** |  |  |  |  |  |  |  |  |
|  | N | Never | Once | Occasionally | Sometimes | Frequently | Almost always | Not applicable |
| 37-Refused to share data with legitimate colleagues | 564 | 535 (94.9) | 11 (2.0) | 6 (1.1) | 0 (0.0) | 2 (0.4) | 2 (0.4) | 8 (1.4) |
| 38-Added one or more authors to a paper who did not qualify for authorship (so-called "honorary authorship") | 563 | 219 (38.9) | 91 (16.2) | 163 (29.0) | 54 (9.6) | 28 (5.0) | 5 (0.9) | 3 (0.5) |
| 39-Accepted authorship for which you did not qualify (so-called "honorary authorship") | 564 | 433 (76.8) | 78 (13.8) | 43 (7.6) | 6 (1.1) | 1 (0.2) | 0 (0.0) | 3 (0.5) |
| 40-Demanded authorship for which you did not qualify (so-called "honorary authorship") | 565 | 550 (97.3) | 8 (1.4) | 3 (0.5) | 1 (0.2) | 0 (0.0) | 0 (0.0) | 3 (0.5) |
| 41-Omitted a contributor who deserved authorship | 563 | 531 (94.3) | 23 (4.1) | 7 (1.2) | 0 (0.0) | 0 (0.0) | 0 (0.0) | 2 (0.4) |
| 42-Submitted (or re-submitted) a manuscript or grant application without consent from one or more of the authors | 563 | 499 (88.6) | 27 (4.8) | 29 (5.2) | 4 (0.7) | 2 (0.4) | 0 (0.0) | 2 (0.4) |
| 43-Submitted the same manuscript to multiple journals at once (so-called "duplicate" or "double submission") | 564 | 558 (98.9) | 4 (0.7) | 0 (0.0) | 0 (0.0) | 0 (0.0) | 0 (0.0) | 2 (0.4) |

Questionable research practices are listed in the order presented on the survey.
<sup>a</sup> Percentages are calculated using data from individuals who responded to the item (i.e., non-responder data are not included in the denominator).

**Box** The top 10 most frequently reported QRPs among an international sample of 590 health professions education researchers (listed from highest to lowest frequency).

1. Added one or more authors to a paper who did not qualify for authorship (so-called “honorary authorship”)
2. Cited articles and or materials that you have not read
3. Selectively cited certain papers just to please editors or reviewers
4. Inappropriately stored sensitive research data (e.g., data that contains personally identifiable information)
5. Selectively cited your own work just to improve your citation metrics
6. Ignored a colleague’s questionable interpretation of data
7. Collected course or curriculum data under the guise of “program evaluation” without human-subjects ethics (IRB) approval with the ultimate intent of using the data for research purposes
8. Inappropriately emailed sensitive research data (e.g., data that contains personally identifiable information)
9. Accepted authorship for which you did not qualify (so-called “honorary authorship”)
10. Spread study results over more papers than is appropriate (so-called “salami slicing”)

## Discussion

This study examined the frequency of self-reported QRPs among HPE researchers, practices that may be detrimental to scientific inquiry.^1,2^ To our knowledge, this is the first study to explore QRPs across the HPE research continuum. Taken together, our findings indicate that a substantial proportion of HPE researchers admit to having engaged in a range of QRPs. These results are consistent with the extant literature on research misconduct in other fields,^4–6,8,16^ and they are important because QRPs can waste resources, provide an unfair advantage to some researchers over others, and ultimately impede scientific progress. Therefore, this study raises significant concerns about the credibility of HPE research, suggesting that our community may need to take a hard look at its ethical norms and research culture.

In our survey, we asked respondents about a range of problematic behaviors, from clear misconduct (data fabrication) and falsification (distortion of results) to plagiarism (copying ideas or words without attribution) and authorship manipulation (e.g., honorary authorship). As Fanelli^8^ noted in his review of research misconduct, certain activities (e.g., honorary authorship or excluding study limitations) are qualitatively different than fabrication and falsification because they do not *directly* distort the quality of the science, per se. However, the damage done to the scientific enterprise by these “less severe” and more ambiguous QRPs may be proportionally greater than deliberate misconduct, for the simple reason that such practices occur more frequently. For example, 20.1% of HPE respondents reported one or more instances of “salami slicing.” While some may think of this as a minor offense, the practice fills the literature with more articles than is seemingly necessary.^1,10^ So, not only does this activity unfairly reward authors and waste resources (e.g., editorial time and journal space), it also can inflate the significance of a given finding, which in turn can distort the outcomes of meta-analyses and other types of systematic reviews.^10,23^

It is worth stating that the interpretation of our results is limited by the nature of the survey methodology employed and, in particular, by the threat of nonresponse bias (especially considering the sensitive nature of the topic under study).^5,6^ Therefore, it is reasonable to ask: (1) how reliable are these frequency estimates, and (2) what can they really say about the actual frequency of QRPs among HPE researchers? Although we did not assess score reliability in the present study, we argue here, as others have previously,^5,8,16^ that self-reports of QRPs likely underestimate the real frequency of questionable behaviors. Researchers who have acted unethically are undoubtedly hesitant to reveal such activities in a survey, despite all assurances of anonymity. What is more, the opposite—researchers admitting to unethical or otherwise questionable practices that they did not do—seems unlikely.^8^ Therefore, we speculate that QRPs may be even more widespread in our community than our estimates imply. Nevertheless, rather than establishing an absolute prevalence of QRPs in HPE, we believe these data are better suited for helping the community understand the nature of the most common QRPs and begin finding feasible solutions to improve our research enterprise.

Questionable research practices related to authorship were some of the most frequently reported behaviors in this study, particularly the practice of giving or accepting unwarranted authorship (so-called “honorary authorship”). Honorary authorship is unethical in academic publishing because individuals who have not sufficiently contributed to the work unfairly receive credit as an author and misrepresent their contributions in the scientific literature.^24^ Our findings corroborate the results of a recent survey of established HPE researchers,^13^ and it seems we are not alone in these practices.^24–26^ For example, a 2008 study of six high-impact medical journals found that 17.6% of corresponding authors admitted to including honorary authors.^24^ In a separate survey of radiology researchers, 58.9% of respondents reported that they had written a paper with a co-author whose contributions did not merit authorship.^27^ In some fields, including HPE, this practice has led journals to require authors to sign an author contribution agreement to verify their explicit authorship roles.^28^ The effectiveness of such requirements is unknown and could be a fruitful area for additional research.

Of note in our findings, the frequency of authors giving honorary authorship and those accepting honorary authorship were not equivalent: 60.6% admitted to adding undeserving authors whereas only 22.7% admitted to accepting honorary authorship. This mismatch suggests that HPE researchers may not fully understand authorship criteria. It may also be a concrete example of the so-called “Muhammad Ali effect”—the idea that individuals often see themselves as more likely to perform good acts and less likely to perform bad acts than others.^29^ Regardless of the mechanism, this finding indicates the need for increased author communication and better shared understanding of author roles and responsibilities, such as those set forth by the ICMJE (http://www.icmje.org/recommendations/browse/roles-and-responsibilities/defining-the-role-of-authors-and-contributors.html).

A complete discussion of all the QRPs assessed in this study is outside the scope of this paper. While readers may debate the degree to which some of these behaviors are unethical or otherwise problematic, we should note, as described above, that seemingly minor infractions can have far-reaching negative consequences. For example, by employing QRPs like p-hacking (i.e., manipulating data or analyses until nonsignificant results become significant)^30,31^ or taking advantage of other types of “researcher degrees of freedom,”^32^ scientists can discover “illusory results”^33^ that actually represent artifacts of their study design and analytic approach, as opposed to legitimate findings that can be replicated.^34,35^ Some have argued that such behaviors are the result of individual researchers responding to a set of incentives, the most important of which are rewards for the *quantity* (not the quality) of their publications.^16,36^ A compelling way to view these behaviors is through the evolutionary lens of natural selection. Smaldino and McElreath^37^ made this point, contending that “The persistence of poor methods results partly from incentives that favor them, leading to the natural selection of bad science. This dynamic requires no conscious strategizing—no deliberate cheating nor loafing—by scientists, only that publication is a principal factor for career advancement.” If one accepts this thesis, then the most effective way to improve research practices, and the quality of the corresponding science, is to change incentive structures at the institutional level.^37^

## Limitations and future directions

The current study has limitations. First, nonresponse bias is a legitimate concern, especially considering the modest response rate (28.2%) in the curated sample. That said, recent research suggests that response rate may be a flawed indicator of response quality and representativeness.^38,39^ Moreover, the wave analysis results indicate that nonresponse bias was limited in our sample. Nonetheless, investigators should build on these initial results by examining QRPs among a larger, more global sample of HPE researchers.

A second limitation relates to the inherent challenge of assessing complex, context-specific research behaviors with a survey that requires respondents to self-assess their own practices.^40,41^ Because some of the QRPs on our survey are judgment calls, their evaluation likely requires more detail and nuance than an individual survey item can provide.^40^ So, while the practice of fabricating data is fairly straightforward (and never justified), the same cannot be said for something like inappropriately storing sensitive research data. The latter practice is open to interpretation: what is considered “inappropriate storage” to one researcher might seem perfectly fine to another. We attempted to reduce this type of ambiguity in our survey items by employing a rigorous expert review process, which resulted in several revised (and we believe, improved) survey items. But, ultimately, “survey self-reports can never fully rule out ambiguities in meaning, limitations in autobiographical memory, or motivated biases.”^40^ Thus, future work might apply qualitative research methods to address some of these limitations and further unpack the nature of QRPs in HPE.

Finally, we administered our survey in English and did not ask respondents to focus on a particular time period. These implementation and design choices could have negatively affected data quality. For example, several respondents noted in their written comments that ethical standards related to human-subjects research had evolved over time. Future research could address this problem by limiting the time period that respondents are asked to consider.^5^

## Recommendations for practice

As HPE continues to mature as a field, it is essential that we explicitly confront our obligation to conduct our research in an ethical and responsible manner. Previously, we suggested a number of recommendations to improve practice,^7^ and the findings reported here indicate the time is *now* to implement these and other policy changes. Our recommended approaches included: (1) empowering research mentors as role models, (2) openly airing research dilemmas and infractions, (3) protecting whistleblowers, (4) establishing mechanisms for facilitating responsible research (e.g., creating HPE-specific institutional review boards^42^), and (5) rewarding responsible researchers (e.g., providing grant funding and publication opportunities for replication studies^34^). These are institution-level approaches; they embrace the idea that QRPs are not simply the result of individual researchers acting badly. Instead, there are important contextual factors that influence researcher behavior (e.g., social norms, power disparities, institutional policies, and academic incentives).^7^ Examples of initiatives being tried in other fields and institutions include promotion and tenure guidelines that privilege publication quality over quantity,^43^ study pre-registration plans,^33^ and other open-science practices (e.g., open data and materials sharing).^44^ All of these approaches require study to determine their efficacy in HPE.

## In summary

Cultivating the responsible conduct of research is essential if we are to maintain scientific integrity and engender public confidence in our research.^1,2^ This study raises significant concerns about the credibility of HPE research and presents a somewhat pessimistic picture of our community. We should be clear though—most HPE scientists surveyed did *not* report the vast majority of QRPs. They are presumably doing good science. Nevertheless, we believe reforms are needed. In addition, we recommend future research to monitor QRPs in HPE and evaluate the effectiveness of policies designed to improve the integrity of our research enterprise.

## Acknowledgments

The authors wish to sincerely thank all of the HPE researchers who took the time (and had the courage) to complete our survey.

**Funding/Support**None reported.

**Other disclosures**
None reported.

**Ethical approval**
The Ethical Review Board Committee of the Netherlands Association for Medical Education approved this study (Dossier #937).

## Disclaimer

The views expressed in this article are those of the authors and do not necessarily reflect the official policy or position of the Uniformed Services University of the Health Sciences, the U.S. Navy, the Department of Defense, or the U.S. Government.

Written work prepared by employees of the Federal Government as part of their official duties is, under the U.S. Copyright Act, a “work of the United States Government” for which copyright protection under Title 17 of the United States Code is not available. As such, copyright does not extend to the contributions of employees of the Federal Government.

---

**Figure 1.**
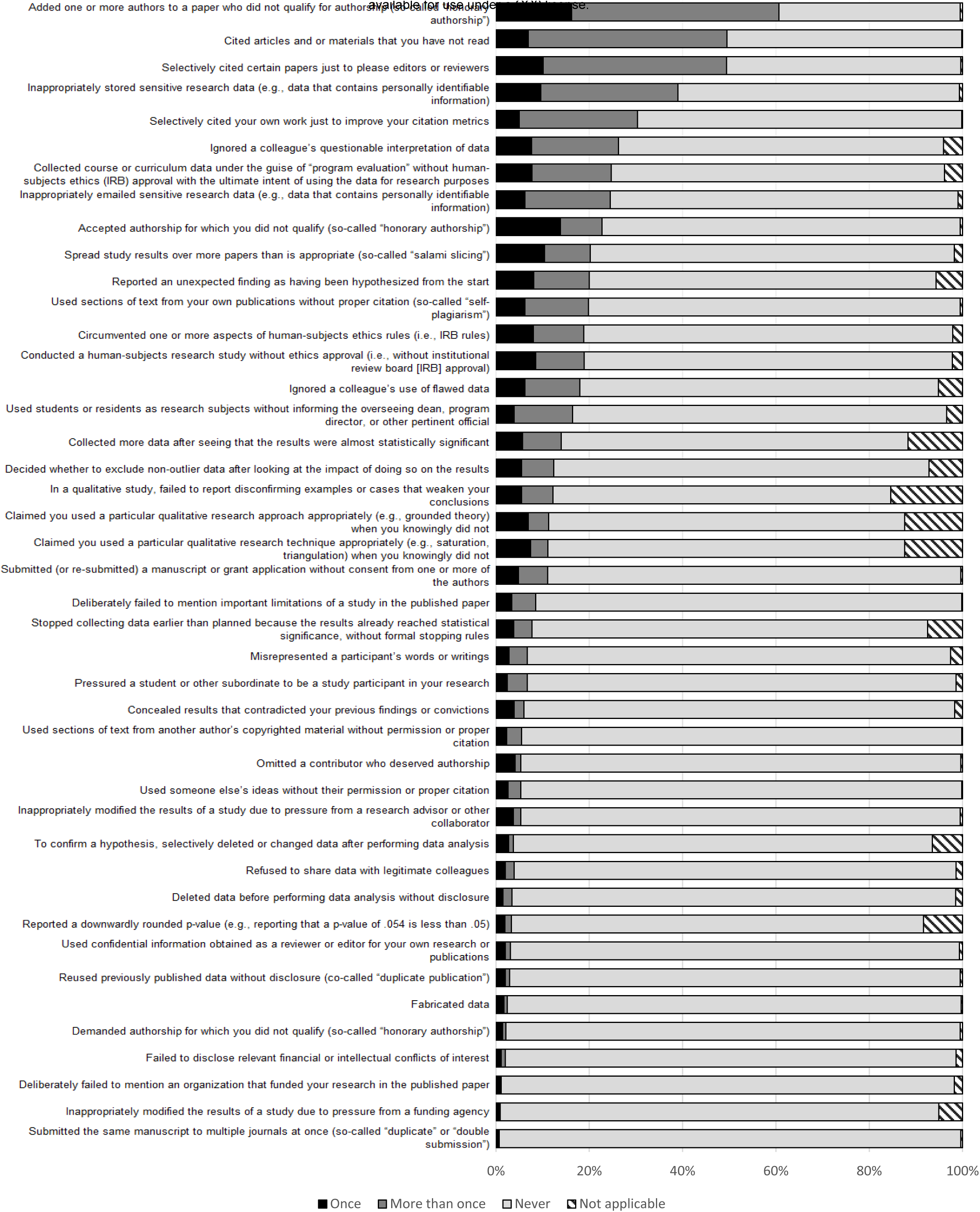
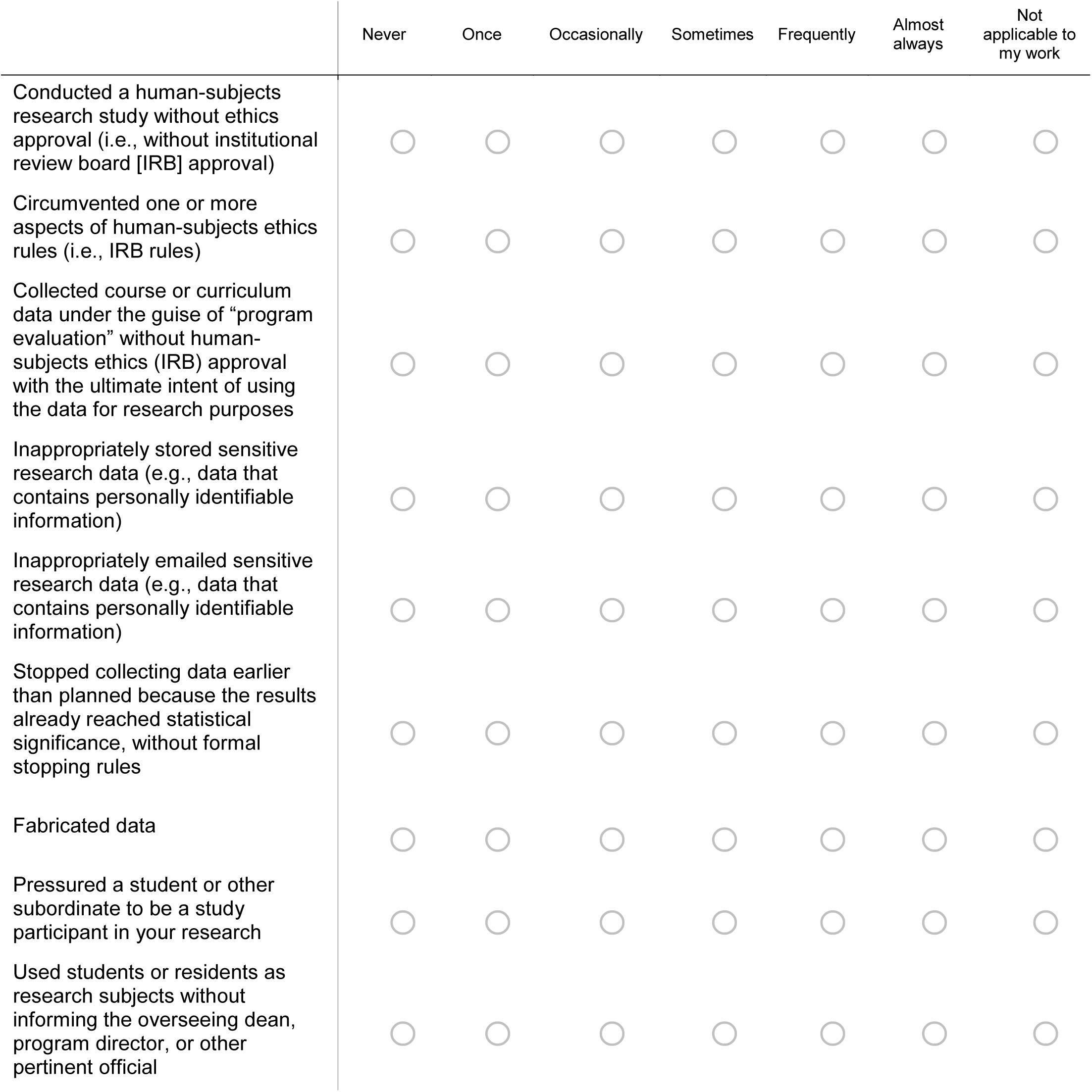
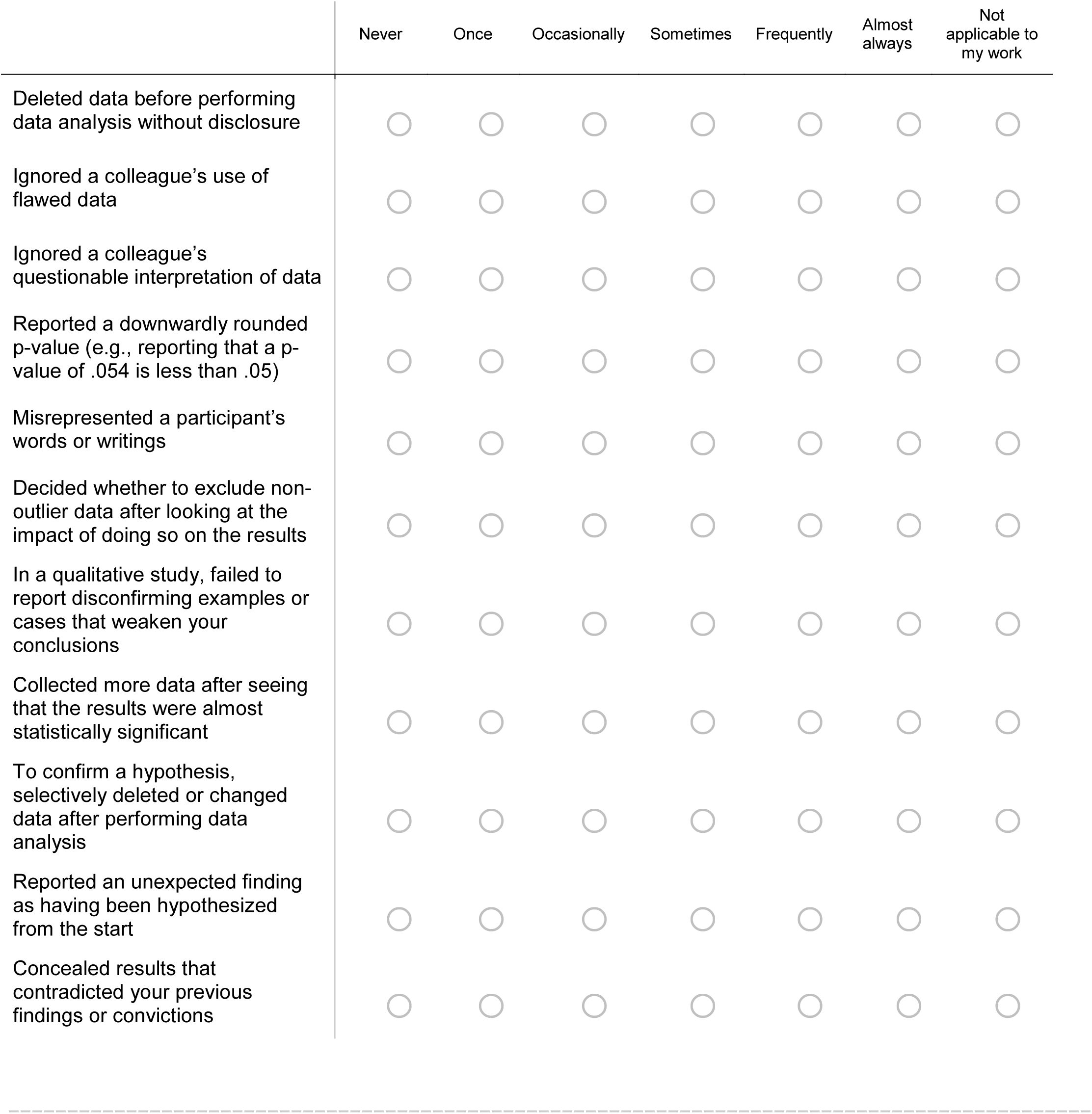
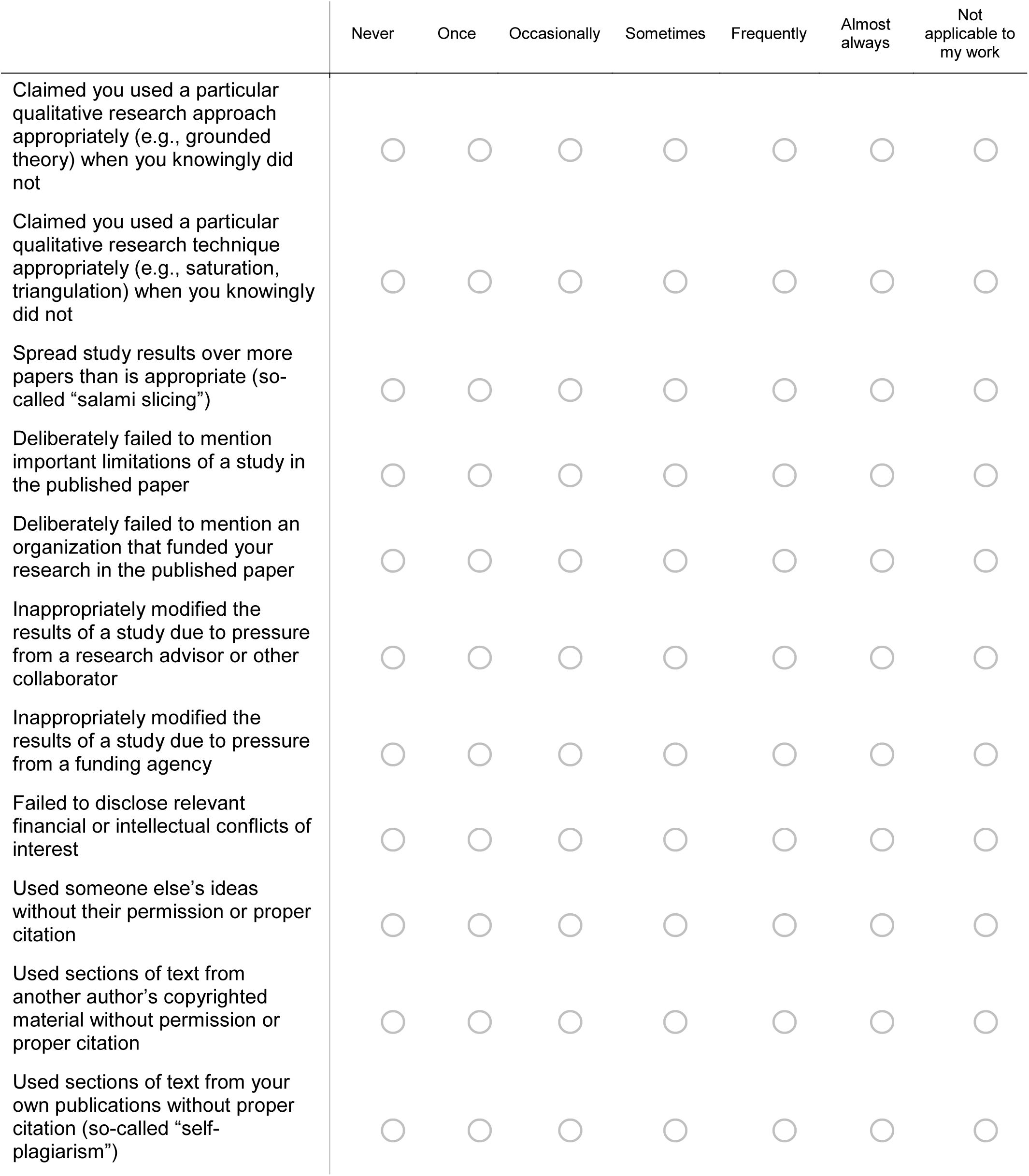
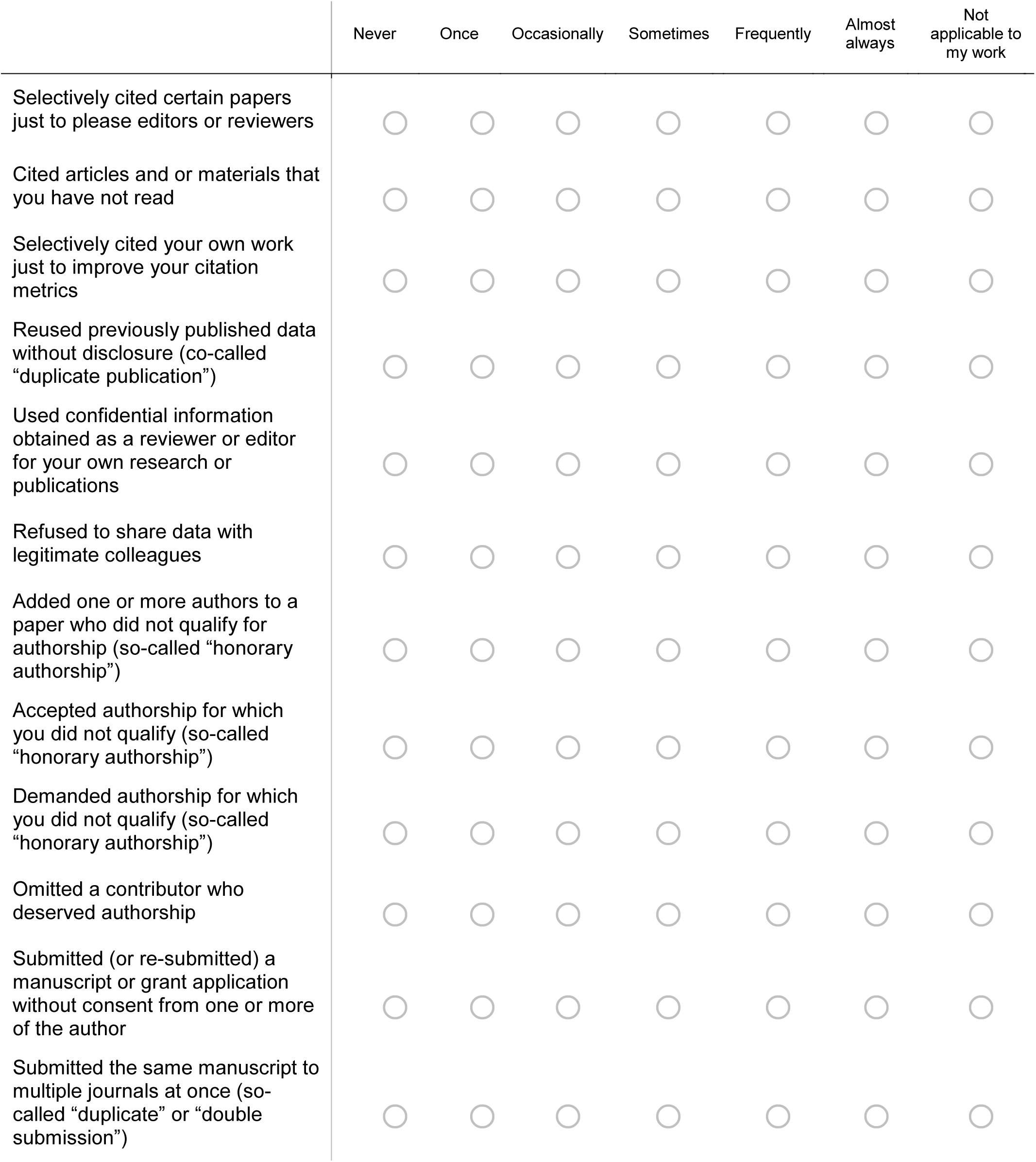
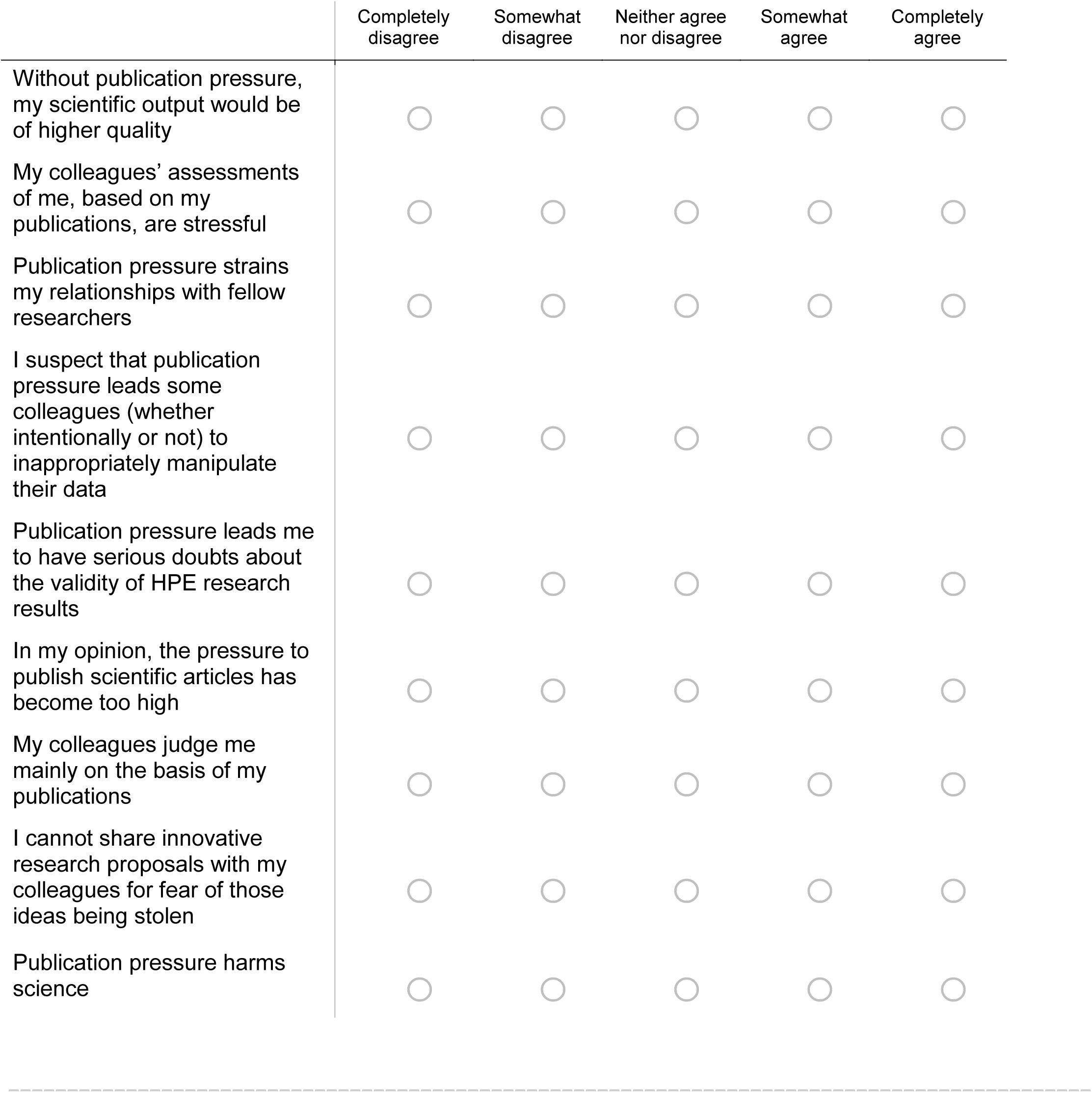
Stacked bar graph showing the frequency of self-reported questionable research practices among an international sample of 590 health professions education researchers.

